# Experimental Control of a Reaction Occurring during the Interaction between Chicken Anemia Virus (CAV) and Its Corresponding Antibodies

**DOI:** 10.1101/2021.02.12.430950

**Authors:** Ognyan Ivanov, Petar Todorov, Konstantin Simeonov, Ashok Vaseashta

## Abstract

In our ongoing investigations, we have studied a specific interaction between electromagnetic fields and matter – the so-called Electromagnetic echo effect (EMEE). It enables rapid and contactless investigations of gases, liquids and solids to be performed, since the signal generated as a result of the effect is quite sensitive to all kinds of changes occurring within the studied samples. The effect can be considered universal for all matter and provides analysis in real time. We use this phenomenon to demonstrate the practical possibility to control reactions, occurring between Chicken anemia virus (CAV) and the corresponding antibodies. This methodology can be used for simple but reliable control of similar, otherwise hard to detect, antigen-antibody reactions, in order to confirm the presence of a certain viral species. The approach offers a high level of safety, since it enables measurements to be taken remotely, thus limiting exposure to contagion. We further discuss the possibility to register the presence of SARS-nCoV-2 in an attempt to address current global pandemic.

## 1. Introduction

A team of researchers from the Bulgarian Academy of Sciences has been actively involved in research of electromagnetic field-matter interactions and development of sensors based on the Electromagnetic echo effect (EMEE) [1], which was previously known as the Surface photo charge effect (SPCE) [2,3]. It was discovered during studies on the transverse acoustoelectric voltage effect and can be observed when any solid body is irradiated by an electromagnetic field. This induces an alternating potential difference between the body and the common electrical ground of the system, which has the same frequency as the frequency of the irradiating field [4]. By measuring the signal from the EMEE, different properties of all kinds of matter can be investigated – solids, liquids and gases. The measurements are almost instantaneous and there is no mandatory need for contact with the studied sample. The devices operating on the basis of the EMEE are cost-effective and easy to operate with, providing convenient and fast way of testing.

## 2. Materials and Methods

Numerous experimental results reveal that the EMEE is induced with frequencies not only in the visible and adjacent regions of the spectrum but also between 1 Hz and 1 GHz. Although no experiments have been done with frequencies between 1 GHz and infrared and higher than ultraviolet, it is expected that the effect is present in the whole electromagnetic spectrum. Additional amplitude modulation is used with emission frequencies in the visible region, because it is still not possible to directly measure signals with frequency higher than the THz range. The EMEE can be distinguished by other similar effects, such as thermal electricity, internal and external photoelectric effect (PEE), etc. Spectral studies show that, for example, unlike the external PEE that has a threshold for the wavelength of irradiation above which it does not exist (~600 nm), the EMEE is still observed [5]. Another difference is that the EMEE exists not only in conductors and semiconductors, but also in dielectrics [3,6,7]. Some hypotheses have been developed [4,8] but there is still no full theoretical explanation for the presence of the effect in all solids.

In conductors, optical radiation falling onto their surface may cause emission of electrons. The theory of the PEE assumes that this effect is caused only by the existence of sharp irregularities on the surface of the conductor, which results in an exponential decay of the wave functions of the electrons [9]. The non-uniform distribution of charges in near proximity to the surface of a conductor forms a so-called double layer at the conductor-vacuum interface. The electrostatic field of the double layer causes free charges to be repelled from the surface and is the main explanation of the PEE in metals.

In the case of the EMEE, the irradiating field causes redistribution of near-surface electrons by creating a force acting on the medium, and thus the electrostatic potential of the double layer changes. In other words, a macroscopic dipole momentum is induced by the falling electromagnetic radiation on the surface of an isolated conductor. If it is connected to another conductor with zero potential, after a certain period of time a flow of electrons will occur between them until a stationary redistribution of the surface charges and potentials is reached [10]. This macroscopic polarization of the conductor can be measured. The change in the potential of the double layer depends both on the surface irregularities and the intensity of the electromagnetic radiation. In conductors, it was experimentally established that the amplitude of the EMEE signal depends on the composition of the material and other parameters, such as the modulation frequency and intensity of the irradiating field [11]. In some cases, even previous illuminations of a sample affects the generated signal [3].

In semiconductors, the incident radiation can induce generation of carriers, if the photon energy is higher than a certain threshold. This value can be, for example, the bandgap energy in the case of electron-hole generation caused by band-to-band excitation. When the intensity is high, the electron-hole density is often also high. As in conductors, the EMEE is generated in semiconductors by the interaction of the free carriers in them and the incident electromagnetic field, but the voltage is caused by the redistribution of charges that are already accumulated in the area close to the surface. Some theories suggest that the photovoltage effect (PVE) and the photo conductivity (PC) mainly contribute to the EMEE in semiconductors [12]. PC is an electrical and optical phenomenon observed when a material becomes more electrically conductive after absorption of electromagnetic radiation. While similar to the PEE, which involves light photons displacing electrons completely out of a material, the PVE involves incident photons dislocating electrons only out of their atomic orbitals but keeping them flowing freely within the material.

In dielectrics, a possible reason for the presence of the EMEE may be dipole molecules present on their surface due to absorption or other phenomena. The force created by the incident electromagnetic radiation is proportional to the gradient of the electrical permittivity of the medium, as in conductors. This leads to redistribution of the absorbing molecules along the surface of the dielectric and thus it becomes electrically charged. It has been found that the generated EMEE voltage depends on the intensity of the field, the type of the irradiated dielectric material and its thickness [7].

There is a possibility that the physical mechanisms of the EMEE may be different for the various types of materials and their explanation may need novel ideas. A possible approach for doing this may be the use of Density functional theory including the effects of external electromagnetic fields.

A basic characteristic of the EMEE is that it strongly depends on the properties of the investigated solids and any changes in them – of the chemical composition, conductivity, surface electrical state, etc. This reveals vast opportunities for studying solids – for example, quality control of raw materials [13,14], semiconductor characterization [12], inspection of mechanical defects, impurities and topology of surfaces [3], and many more. All kinds of fluids can also be investigated by placing them in contact with the solid under irradiation [15]. Any changes in the properties of a fluid, or processes taking place in it, are transferred to the solid-fluid interface and can be thus detected [16,17]. It has been previously demonstrated that the EMEE can be utilized to distinguish between different liquids [18]; to determine the octane factor of gasoline [19], or the quality of drinking water from the public water distribution system [4]; to control fog parameters, including the presence of impurities in fog [20]; and even to monitor processes in biological fluid samples [21].

The general setup for measuring the EMEE signal in the visible range of the spectrum is presented in Figure 1. Most often, a laser module is used as a radiation source (R). The laser beam is modulated by a signal modulator (M) and irradiates the solid sample (S), specially chosen for the specific task. The signal generated in the solid is “captured” by the electrode (E), which can be in near proximity or in contact with the sample, and then transferred to a nanovoltmeter (N) for measurement.

**Figure 1.**
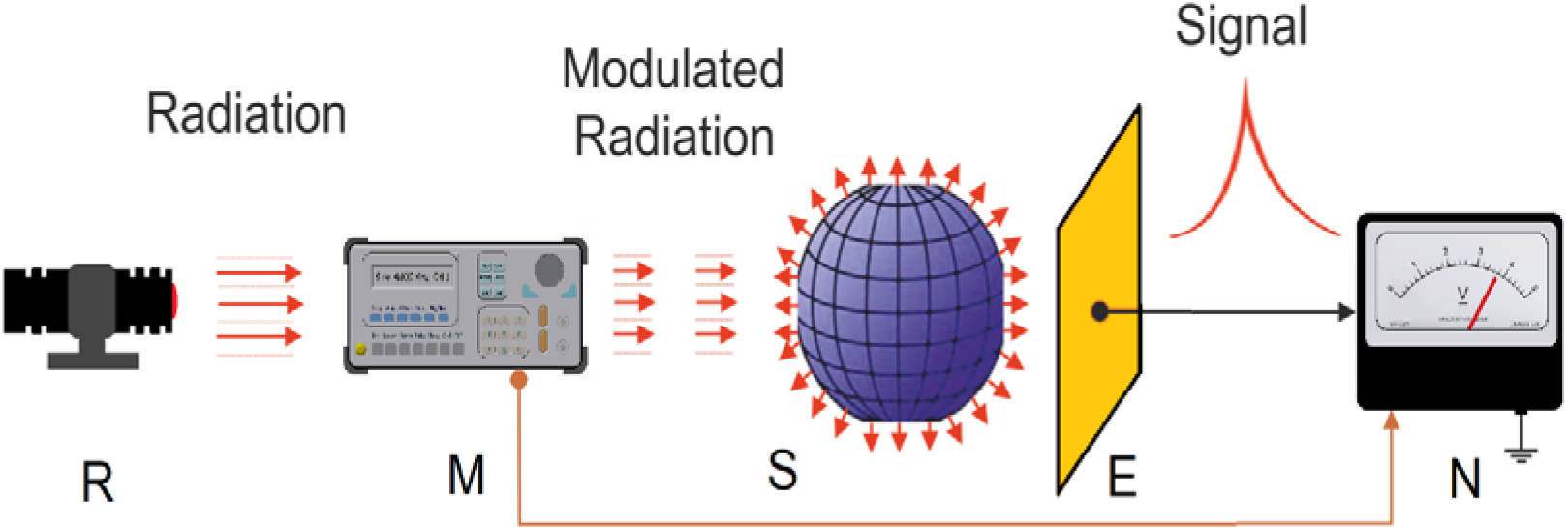
General setup for EMEE measurement: R – source of electromagnetic radiation; M – signal modulator; S – solid sample; E – electrode; N – nanovoltmeter.

Since the signal from the EMEE is very weak (on the micro- and nanovolt scale), it is important to shield the setup from external electromagnetic interferences and to use a nanovoltmeter, which is capable of separating the useful signal from the background noise. This can be achieved by using a phase-sensitive detection and a reference signal from the signal generator.

The concept for experimental investigation of fluids with the EMEE involves a contact between the surface of the irradiated solid and the fluid. Any alterations in the properties of the fluid will cause corresponding changes at the interface (the boundary surface between the solid and the fluid), where the signal is generated from (especially in the irradiated region). In this way, if all other parameters remain the same, the registered variation of the signal is due only to the specific changes in the fluid. For example, if a suitable solid sensing structure is selected, a very small amount of liquid, such as a tiny droplet, is enough to form an interface with the solid to perform analysis [19].

The idea of using the EMEE to detect the presence of viruses, and SARS-nCoV-2 in particular, has already begun to develop [22]. While SARS-nCoV-2 was unknown to medical science until late 2019, the family of viruses are not new. SARS-nCoV-2 is an enveloped, positive-sense, single-stranded RNA beta coronavirus of approximately 30 kb in length. The RNA has a 5’-cap and a 3’-poly(A) tail and can act as an mRNA for immediate translation of the viral polyproteins. In addition, both 5’- and 3’-ends of the RNA present a highly structured untranslated region (UTR) that plays an important role in the regulation of RNA replication and transcription. SARS-nCoV-2 gets into the cell through recognition by the spike glycoprotein present on the surface of the virus envelope of the angiotensin converting enzyme 2 (ACE2) receptors. Once inside the cell, the infecting RNA acts as a messenger RNA (mRNA), which is then translated by host ribosomes to produce the viral replicative enzymes, which generate new RNA genomes and the mRNAs for the synthesis of the components necessary to assemble the new viral particles.

Based on the long experience of our team with creation of various sensors, this method promises to be quite useful for rapid identification of viral threats, if elaborated further. Initial experiments with avian coronavirus causing infectious bronchitis (strain Massachusetts) confirm the reliability of the approach [23]. By irradiating a thin strip layer of a virus solution on the surface of a solid substrate, a certain EMEE signal is generated. When adding a micro-droplet of an antibody-containing serum, a specific antigen-antibody reaction occurs and rapidly changes the amplitude of the output signal. In this way, the interaction is confirmed. The experiments conducted so far show that it is possible to control the presence of viruses by using this technique. Since antibodies are more expensive than viruses, we detect the specific reaction by adding a small droplet of the antibodies-containing serum to a layer containing the virus. When developing a real-life functioning sensor for this purpose, the setup will be reversed – the antiviral antibodies will be immobilized on the solid surface, and in case the corresponding virus comes in contact with it, the reaction between them will be registered, thus alarming about the presence of the virus. There is a possibility to make a solid layer containing the antibodies, in order the sensor to be able to be used for numerous detections. Here we present the principle used for detection, so that the experiments can be replicated by anyone, but the effectiveness of the system is determined by setting specific technological parameters, which is know-how of our team. Such parameters include the type of the structure generating the signal, the wavelength and intensity of the incident radiation, the measurement conditions and technique, etc.

## 3. Results

Here we present experiments with Chicken anemia virus (CAV) and the corresponding antibodies. This virus was chosen because of its importance as an economically significant pathogen for industrial poultry production. Clinically manifested CAV infections are relatively rare and occur in chickens between 14 and 21 days of age; however, the economic significance of this virus is due mainly to its immunosuppressive potential. Birds of different ages could be infected and develop an immune deficiency, resulting in exalted secondary or concomitant infections, whose clinical manifestations prevail over the CAV related signs, thus the role of CAV as a trigger for these conditions could remain hidden. In addition, CAV was detected in SPF (specific-pathogen-free) flocks, widely used in research as well as a source of eggs used for vaccine production [24]. In recent years, there have been a significant number of publications concerning the unwanted CAV contamination of attenuated vaccines against other viral pathogens produced in such eggs [25,26]. The use of a vaccine contaminated with exogenous virus would lead to an outbreak of the disease, or at least to spreading of the infection among flocks, in which these vaccines are administered [27]. Therefore, the CAV detection is included among the list of regular monitoring procedures for exogenous viral contamination of attenuated poultry vaccines.

At present, molecular detection methods, such as PCR (polymerase chain reaction), are used to detect CIAV (chicken infectious anemia virus). However, some types of PCR such as the conventional PCR have been shown to be insufficiently sensitive due to the low dose of CIAV in live vaccines [28]; it is therefore necessary to look for other means of detection. In this study, we use the interaction between CAV and its corresponding antibodies as a model to show the applicability of the EMEE-based sensing technique for control of the emergence of different viral species. The setup is very similar to the one used for the infectious bronchitis (strain Massachusetts) described above [23], and is shown in Figure 2.

**Figure 2.**
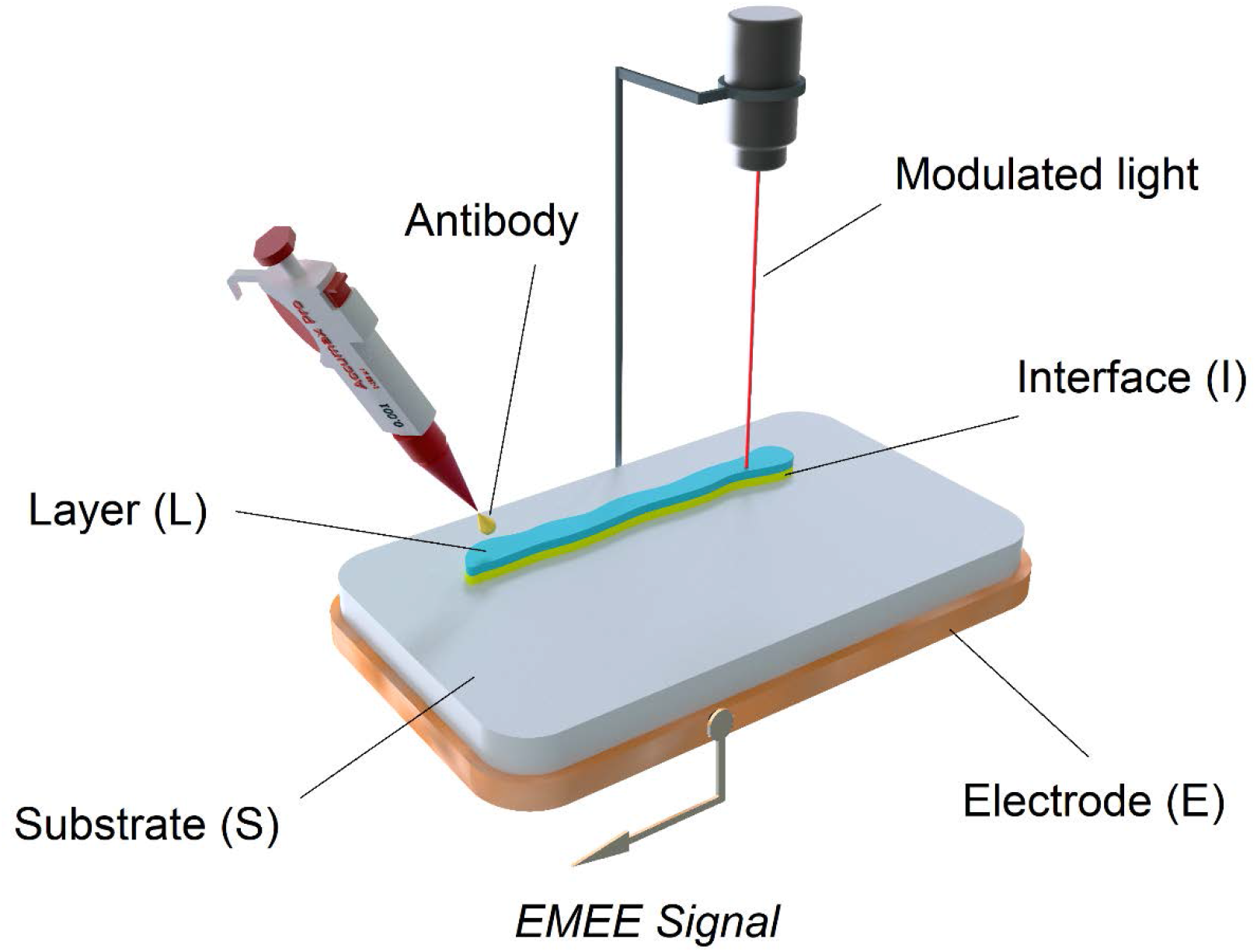
An EMEE setup for CAV-antibody reaction detection: S – solid substrate; L – contact layer containing CAV; I – solid-liquid interface, generating the signal; E – measuring electrode.

The Nobilis CAV®P4 vaccine was used as a source of virus in all experiments. The vaccine, containing live CAV virus strain 26P4 was diluted in Unisolve diluent according to manufacturer’s recommendations. Briefly, the freeze-dried pellet was reconstituted in 13 ml diluent, thus every 13 μl containing at least 10^3^.TCID50 of virus. To avoid repeated freezing-thawing of once diluted vaccine, it was distributed in vials (300 μl/vial) and stored at −20° C until use. Positive and negative controls from a diagnostic kit IDEXX CAV Ab Test (IDEXX Laboratories, Netherland) were used as antibodies-containing and containing no antibodies sera, respectively. The latter were used undiluted in the experiments.

A thin strip-like layer (L) of a solution containing CAV (about 15 x 2 mm in length and width, correspondingly) was spread on the solid substrate of the sensing structure, thus forming an interface between them. The layer was irradiated with a modulated laser light at one end, which induced a corresponding EMEE signal, sensed by the electrode (E). The initial value of the signal remained stable at around 5 relative units (Figure 3). At a certain moment, a tiny droplet containing antibodies for CAV was applied at the other end of the layer, in order to avoid interfering with the laser beam and, thus, falsely changing the signal. The addition of antibodies almost instantly produced a sharp peak in the measured signal with a value of about 16 relative units, after which the amplitude returned to the initial value. This strong deviation (~320%) of the amplitude of the signal proves that the antigen-antibody reaction has been registered.

**Figure 3.**
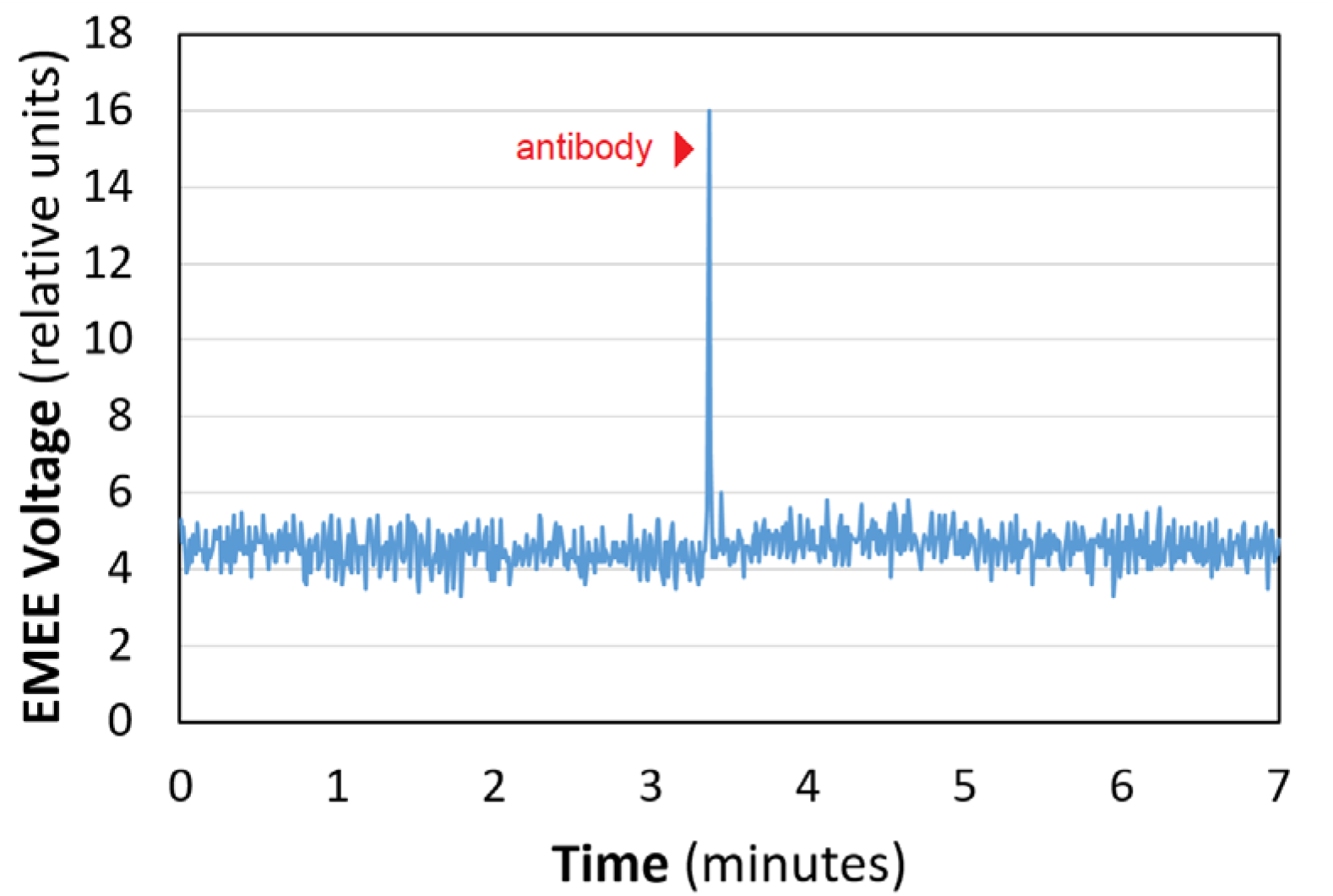
Recording of the EMEE signal variation. The layer on the sensor surface contains the CAV. An arrow indicates the moment when a 10 μl sample from the antibody-containing serum is added.

When repeating the experiment with a control serum containing no specific antibodies (Figure 4), soon after the micro-droplet was added, there was only a slight change in the signal – from the initial value of about 6 relative units to a small peak of almost 9 relative units, after which the signal returns to about the initial value. Here, the small deviation (less than 50%) was due to the change of the composition of the layer.

**Figure 4.**
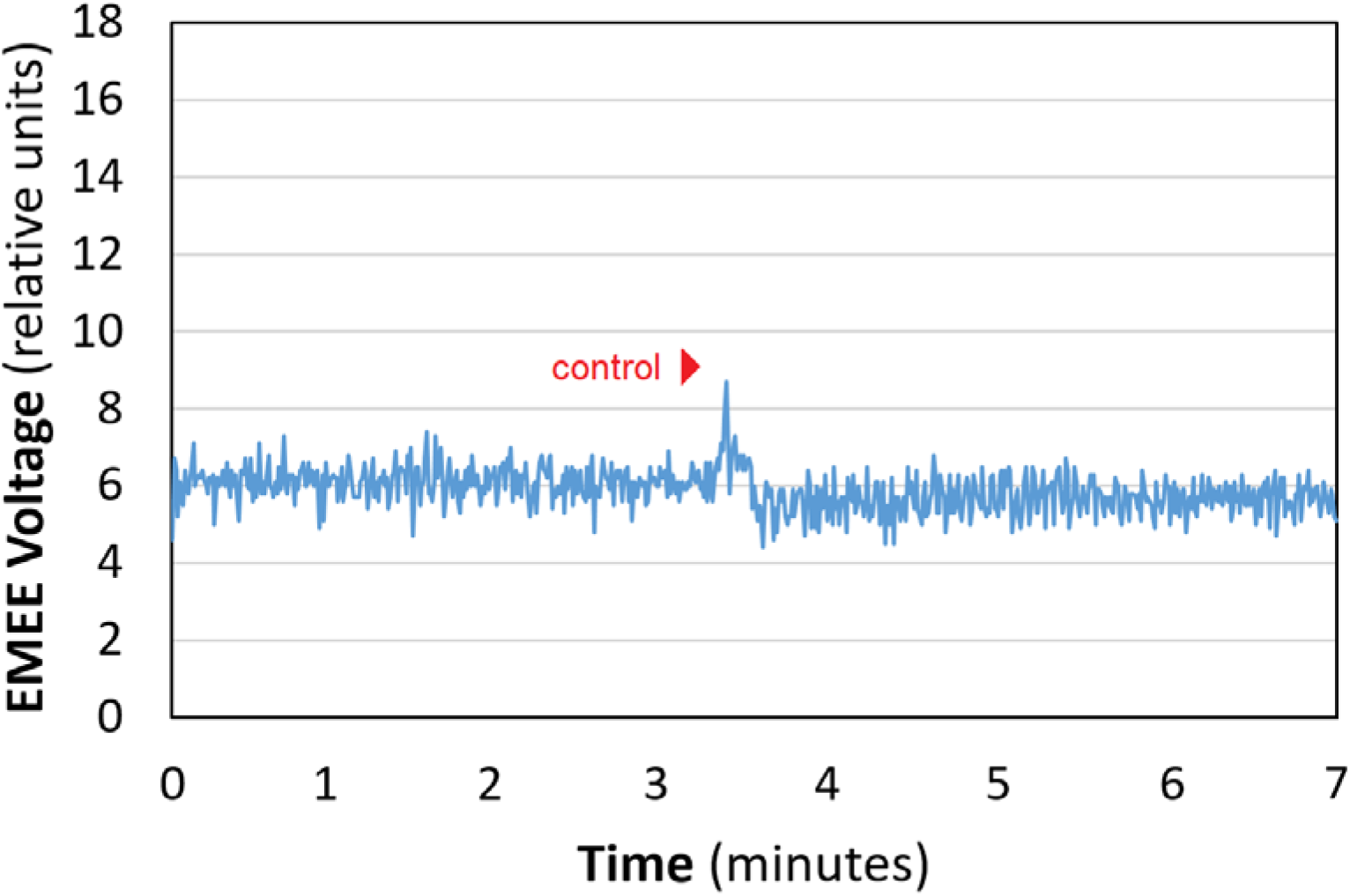
Recording of the EMEE signal variation. The layer on the sensor surface contains the CAV. An arrow indicates the moment when a 10 μl sample from the control serum is added.

The amplitude of the signal in Figure 3 and Figure 4 is given in relative units, because the measuring equipment transforms the real values. Nevertheless, the proportions between the values are kept the same, which is enough to make comparisons. In both figures, the signal was recorded at an interval of 0.01 min.

## 4. Discussion

The experiments presented here demonstrate that sensors operating on the basis of the EMEE can be developed to support the already known means of different viral species detection. The quick chemical interaction between a CAV with a serum containing its corresponding antibodies was registered almost instantly upon contact. This produced a very strong and sharp peak in the measured amplitude of the EMEE signal. In contrast, when the virus was exposed to a control serum, only a small deviation (more than six times smaller) of the signal was observed, meaning that a clear distinction can be made between the two cases. Moreover, the results demonstrate that very small amounts of reagents are enough to induce a measurable difference in the signal. This suggests that such devices can be successfully developed to reliably detect the presence of SARS-CoV-2, which will be our next goal. The initial investigations that support this idea have already shown promising results in the case of detection of an avian infectious bronchitis virus, where similar results were obtained [23]. The main advantage of the demonstrated method is that it can be applied for precise, rapid and cost-effective monitoring of indoor air, inspection of surfaces or human body fluids for the presence of a certain virus, since the EMEE response is strictly specific. Similar sensing devices have been already developed for detection of harmful air pollution in the form of dispersed CBRN (chemical, biological, radiological and nuclear) agents, as well as for tracking changes in the chemical composition and microphysical properties of fog [16], under an FP-7 Security Programme project. Results of previous experiments suggest that the same methodology can also be used for quick and contactless evaluation of the authenticity of virus vaccines – whether they have been stored properly or have expired due to changes in the substance [14].

## 5. Conclusions

A novel approach for viral detection utilizing EMEE sensing has been presented. Experimental results regarding control of the presence of a Chicken anemia virus by registering the specific reaction with its corresponding antibodies prove the feasibility of the proposed method. The development of a fully functioning sensing system that can provide almost instantaneous results is crucial for effective health protection and quick reaction against viral outbreaks, such as the current coronavirus pandemic. The devices operating on the basis of the EMEE are capable of fulfilling these needs. In addition, they have low cost and maintenance requirements, offer fast and precise measurement, and their size can be minimized, so that portability can be ensured.

## Author Contributions

Conceptualization, O.I.; methodology, O. I.; investigation, O. I. and P.T.; formal analysis, O. I. and A.V.; visualization, P.T.; writing—original draft preparation, P.T. and K.S.; writing—review and editing, K.S. and A.V.; supervision, O.I. and A.V. All authors have read and agreed to the published version of the manuscript.

## Institutional Review Board Statement

Not applicable.

## Informed Consent Statement

Not applicable.

## Data Availability Statement

Data is contained within the article.

